# A high-density narrow-field inhibitory retinal interneuron with direct coupling to Müller glia

**DOI:** 10.1101/2020.01.23.917096

**Authors:** William N Grimes, Didem Göz Aytürk, Mrinalini Hoon, Takeshi Yoshimatsu, Clare Gamlin, Daniel Carrera, Richard M Ahlquist, Adit Sabnis, Jeffrey S Diamond, Rachel O. Wong, Connie Cepko, Fred Rieke

**Author notes:** Allen Institute for Brain Science, Seattle, WA. These authors contributed equally (William N Grimes, Didem Göz Aytürk, Mrinalini Hoon).

## Abstract

Amacrine cells are interneurons comprising the most diverse cell type in the mammalian retina. They help encode visual features such as edges or directed motion by mediating excitatory and inhibitory interactions between input (i.e. bipolar) and output (i.e. ganglion) neurons in the inner plexiform layer (IPL). Like other brain regions, the retina also contains glial cells that contribute to neurotransmitter uptake, neurovascular control and metabolic regulation. Here, we report that a previously poorly characterized, but relatively abundant, inhibitory amacrine cell type in the mouse retina is coupled directly to Müller glia. Electron microscopic reconstructions of this amacrine type revealed extensive associations with Müller glia, whose processes often completely ensheathe the neurites of this amacrine cell type. Microinjections of small tracer molecules into the somas of these amacrine cells led to selective labelling of nearby Müller glia, leading us to suggest the name “Müller glia-coupled amacrine cell” or MAC. Our electrophysiological data also indicate that MACs release glycine at conventional chemical synapses with amacrine, bipolar and retinal ganglion cells (RGCs), and viral transsynaptic tracing showed connections to several known RGC types. Visually-evoked responses revealed a strong preference for light increments; these “ON” responses were primarily mediated by excitatory chemical synaptic input and direct electrical coupling to other cells. This initial characterization of the MAC provides the first evidence for neuron-glia coupling in the mammalian retina and identifies the MAC as a potential link between inhibitory processing and glial function.

**Significance Statement:** Gap junctions between pairs of neurons or glial cells are commonly found throughout the nervous system, and play a myriad of roles including electrical coupling and metabolic exchange. In contrast, gap junctions between neurons and glia cells are rare and poorly understood. Here we report the first evidence for neuron-glia coupling in the mammalian retina, specifically between an abundant (but previously unstudied) inhibitory interneuron and Müller glia.

## Introduction

The circuitry of the mammalian retina employs diverse, parallel circuits to extract many distinct features of the visual world. Amacrine cells, the largest class of inhibitory cell types in the retina, play a critical role in such processing. Amacrine cells are interneurons with cell bodies in either the inner nuclear layer (INL) or ganglion cell layer (GCL), and their processes are typically confined to the most extensive synaptic region of the retina, the inner plexiform layer (IPL; Figure 1). They employ a diverse array of synaptic mechanisms, including the release of many different neurotransmitters (e.g. GABA, glycine, glutamate, acetylcholine, nitric oxide, dopamine, neuropeptides), as well as gap junctions (i.e. electrical synapses; for review see (Kolb, 1995, Masland, 2012, Wu and Maple, 1998)). Understanding the connectivity and function of amacrine cells is essential to our understanding of how the retina performs computations required for effective and rapid visual processing.

**Figure 1.**
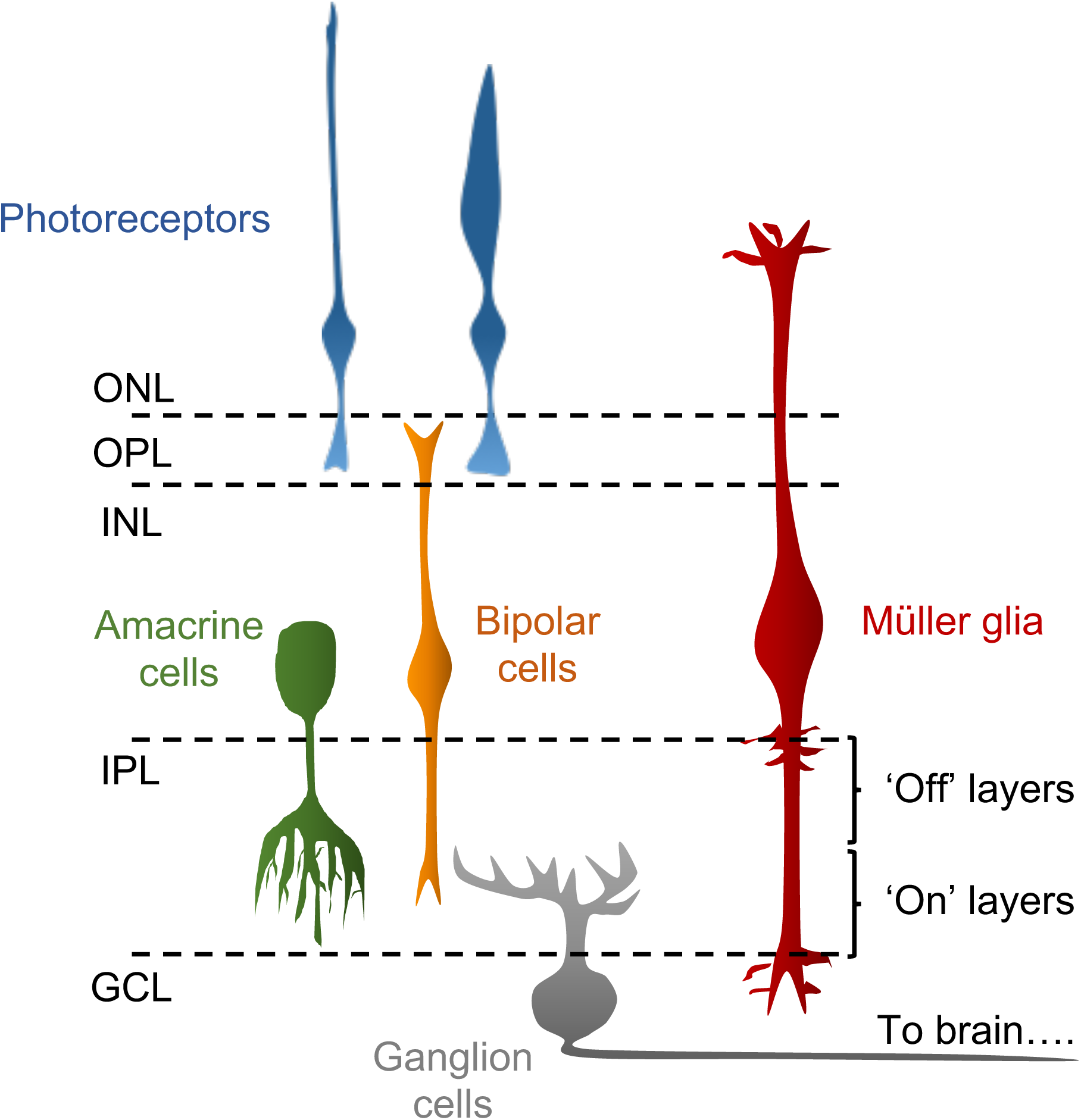
Retinal Schematic. Light is captured by rod and cone photoreceptors and converted into neural signals. Light-evoked signals are conveyed from photoreceptors to bipolar cells through glutamatergic synapses in the outer plexiform layer (OPL). Bipolar cells relay signals to amacrine and ganglion cells through glutamatergic synapses in the inner plexiform layer (IPL). Synaptic connections between bipolar, amacrine and ganglion cells in the IPL play an important role in feature extraction and the decorrelation of retinal output. Müller glia support retinal function. GCL Ganglion cell layer. INL Inner nuclear layer. ONL Outer nuclear layer.

Here, we examine physiological and anatomical characteristics of a previously unstudied, high-density narrow-field amacrine cell. In addition to presenting cell-specific tools, and describing the cell’s synaptic connections with bipolar, amacrine and ganglion cells, we provide evidence for previously unknown, direct coupling between a retinal neuron and Müller glia. Electron microscopic reconstructions of this amacrine revealed that Müller glia extend processes that physically ensheathe this amacrine cell type’s neurites in the IPL. Small tracer molecules injected into this amacrine cell diffused into nearby Müller glia, indicating gap junction connections between the two cell types and leading us to suggest the name “Müller glial-coupled amacrine cell” (MAC) for this amacrine cell type. Taken together, the results presented here suggest that MACs provide local glycinergic inhibition within retinal neural circuits and also mediate interactions with glial cells.

## Materials and Methods

### Electrophysiology

WT and genetically-modified (see Table 1) C57/Bl6 mice of either sex were used in accordance with Institutional Animal Care and Use Committee at the University of Washington, and were dark-adapted overnight prior to cervical dislocation. Eyes were then immediately enucleated and submerged in warm (∼32° C) bicarbonate Ames medium (Sigma) that was continuously bubbled with carbogen (95% O_2_/5% CO_2_). Small scissors were used to remove the cornea. The lens and vitreous were then removed with forceps before storing the retinal cups for up to 6 hours in a customized dark ‘puck’ containing warm oxygenated Ames. To prepare the retina for electrophysiology recordings, we took half an eye cup, isolated the retina from the pigment epithelium, trimmed the retina into a rectangle and mounted it photoreceptor-side down on poly-L-lysine coated microscope slide. A mounted retina was placed in the recording chamber where it was continuously superfused with warm oxygenated Ames. For experiments that probed visual responses retinal dissections and mounting were conducted exclusively under infrared illumination.

**Table 1.** Mouse lines used in the MAC study

| **Mouse line** | **Driver gene** | **Retinal cells labeled** | **Source** |
| --- | --- | --- | --- |
| *Dact2-eGFP* | Dapper Antagonist of Catenin 2 | Subset of MACs | MMRRC (Stock # 034338-UCD) |
| *Plcxd2-eGFP* | Phosphatidylinositol-specific phospholipase C, X domain containing 2 | Subset of MACs, other amacrine | MMRRC (Stock # 034741-UCD) |
| *NEX-Cre(Ai9)* | Neurogenic differentiation 6 | Subset of MACs, other ACs, horizontal cells | Goebbels et al., 2006 |
| *Dact2-Cre(Ai27)* | Dapper Antagonist of Catenin 2 | Subset of MACs, unknown wide-field AC, unknown RGCs | MMRRC (Stock # 036640-UCD) |
| *PSD95-creEnabled (mVenus)* | Post-synaptic density protein 95 | RGCs, ACs, horizontal cells, Off bipolar cells | The Jackson Laboratory (Stock # 029242) |
| *VGLUT3-Cre(Ai9)* | Vesicular glutamate transporter 3 | RGCs, VGLUT3+ ACs | Grimes et al., 2011 |

Voltage-clamp whole-cell recordings were conducted with electrodes (RGC: 2-3 MΩ, ACs: 5-6 MΩ) containing (in mM): 105 Cs methanesulfonate, 10 TEA-Cl, 20 HEPES, 10 EGTA, 2 QX-314, 5 Mg-ATP, 0.5 Tris-GTP and 0.1 Alexa (594) hydrazide (∼280 mOsm; pH ∼7.3 with CsOH). Current-clamp whole-cell recordings were conducted with electrodes (ACs: 5-6 MΩ) containing (in mM): 123 K-aspartate, 10 KCl, 10 HEPES, 1 MgCl_2_, 1 CaCl 2, 2 EGTA, 4 Mg-ATP, 0.5 Tris-GTP and 0.1 Alexa (594) hydrazide (∼280 mOsm; pH ∼7.2 with KOH). NBQX (10 μM; Tocris) and APV (25 μM) were added to the perfusion solution as indicated in Figure 5. APB (10 μM), UBP (25 μM), APV (25 μM) and NBQX (10 μM) were bath applied for >3 minutes before testing for ChR2-evoked responses in RGCs. To isolate excitatory or inhibitory synaptic input, cells were held at the estimated reversal potential for inhibitory or excitatory input of ∼−60 mV and ∼+10 mV. Absolute voltage values were corrected for liquid junction potentials (K + -based = −10.8 mV; Cs + -based = −8.5 mV).

Full field illumination (diameter: 500-560 μm) was delivered to the preparation through a customized condenser from short wavelength (peak power at 405 or 460 nm) LEDs. Light intensities (photons/μm^2^/s) were converted to photoisomerization rates (R*/photoreceptor/s) using the estimated collecting area of rods and cones (0.37 μm^2^ and 0.5 μm^2^, respectively), the LED emission spectra and the photoreceptor absorption spectra ((Govardovskii et al., 2000)). Flashes were 10 ms in duration, except for ChR2 experiments which utilized a 50 ms flash. In Figure 5A-B the video monitor was set to a mean of ∼200 R*/rod/s.

Electrophysiology example traces presented throughout the figures represent the average of 5-20 raw responses to the same stimuli. All data are presented as mean ± SEM and two-tailed paired Student’s t-tests were used to test significance.

### Immunohistochemistry

For immunohistochemical experiments, retinae were isolated in cold oxygenated mouse artificial cerebrospinal fluid (ACSF, pH 7.4) containing (in mM): 119 NaCl, 2.5 KCl, 2.5 CaCl2, 1.3 MgCl2, 1 NaH2PO4, 11 glucose, and 20 HEPES. Retinas were fixed in 4% paraformaldehyde in ACSF for 20 mins, washed in 0.1M phosphate buffered saline (PBS) and were either incubated with primary antibodies in blocking solution (5% donkey serum and 0.5% Triton in PBS) or were embedded in 4% agar (low-gelling agarose, Sigma Aldrich) and sectioned at 120 μm in the vibratome. Retina slices were utilized for performing immunolabeling with the glycine antibody (gift from D. Pow). GFP was amplified with an anti-chicken GFP antibody from Abcam and retina slices were incubated overnight with primary antibodies in blocking solution followed by washes in PBS and a 3-hour incubation with anti-isotypic secondary antibodies (Invitrogen or Jackson Immunoresearch). Labeling of *Dact2-GFP* or *Plcxd2-GFP* retinas for GAD67 (mouse monoclonal, Millipore), glycine transporter GlyT1 (goat, Millipore), GlyRα1 (mouse, Synaptic systems), syntaxin 1 (mouse, Sigma-Aldrich), and PPP1R17 (rabbit, Sigma-Aldrich) was performed by incubating whole retinas over 3 nights in primary antibody solution followed by washes in PBS and an overnight incubation with anti-isotypic fluorescently conjugated secondary antibodies.

RGCs were biolistically labeled using the CMV:tdTomato plasmid as described previously (Morgan et al., 2008). For tracer injection experiments, 4% Neurobiotin or 4% Neurobiotin and 2% Lucifer yellow (in 200 mM KCl) were injected into GFP positive amacrine cells in the *Plcxd2-GFP* or *Plcxd2-GFP:PSD-95-YFP* lines. Retinas were subsequently fixed for 20 mins in 4% paraformaldehyde in ACSF. To amplify the tracers post-fixation, streptavidin conjugated to Alexa Fluor (Invitrogen) was used for Neurobiotin and an anti-Lucifer yellow antibody (rabbit, Invitrogen) was used for Lucifer yellow. To label postsynaptic glycine receptors, Glycine receptor alpha1 specific monoclonal antibody from Synaptic Systems was utilized and whole mount labeling were performed according to the procedure described above. To label VGluT3 amacrine cells, we used a VGluT3 antibody from Synaptic Systems (Cat. # 135 203). All images were acquired with an Olympus FV1000 microscope using a 1.35 NA 60X oil objective, and the software Amira (Thermo Fisher Scientific) was utilized for visualization and 3D-rendering. To visualize synaptic contacts within RGCs, the RGC processes were first masked in 3D using the *Labelfield* function in Amira. Thereafter, the presynaptic amacrine GFP signal was multiplied with the RGC cell mask (using the *Arithmetic* function in Amira) to isolate the points of apposition between the GFP amacrine processes and RGCs. The immunolabeled glycine receptor signal was also multiplied with the RGC 3D mask to isolate the receptor signal exclusively within the RGC processes. Thereafter, the GFP signal apposed to RGC processes and the glycine receptor signal within RGC processes was co-visualized in 3D to ascertain synaptic appositions.

### Near-infrared branding (NIRB) and serial block face scanning electron microscopy (SBFSEM)

Mouse retina from *Dact2-GFP* transgenic line was flat-mounted on a nitrocellulose filter paper and fixed in 4% glutaraldehyde in 0.1M sodium cacodylate buffer (pH7.4) for 20 min. The tissue was then washed three times and cover slipped in 0.1 M sodium cacodylate buffer for multiphoton imaging. MACs were identified based on their dendrite stratification level. An image stack was taken using a custom-built two-photon microscope with a Ti:sapphire laser (Spectra-Physics) with voxel dimensions of the multiphoton image stack were xy: 0.15 µm and z-step: 0.5 µm. To relocate the MAC using EM, fiduciary marks were burned into the tissue using the near-infrared branding (NIRB) method (Bishop et al., 2011, Della Santina et al., 2016). The laser power was typically set at 100 mW measured at the light path before the objective lens. Repeated line scans for a total duration of 10 s at a wavelength of 780 nm were performed; the laser power was increased by 20 mW at a time if burning was not apparent from the autofluorescence. A box was branded at two different tissue depths; at the cell soma and at the RGC layer. After NIRB-ing, the retina was unmounted and fixed in 4% glutaraldehyde overnight, and then processed for SBFSEM. The tissue was washed 3 x 5 min (all washes) in 0.1M sodium cacodylate buffer and incubated in a solution containing 1.5% potassium ferrocyanide and 2% osmium tetroxide (OsO4) in 0.1M cacodylate buffer (0.66% lead in 0.03M aspartic acid, pH 5.5) for 1 hour. After washing, the tissue was placed in a freshly made thiocarbohydrazide solution (0.1g TCH in 10 ml double-distilled H_2_0 heated to 600 C for 1 h) for 20 min at room temperature (RT). After another rinse, at RT, the tissue was incubated in 2% OsO4 for 30 min at RT. The samples were rinsed again and stained *en bloc* in 1% uranyl acetate overnight at 4° C, washed and stained with Walton’s lead aspartate for 30 min. After a final wash, the retinal pieces were dehydrated in a graded ice-cold alcohol series, and placed in propylene oxide at RT for 10 min. Finally, the samples were embedded in Durcupan resin. Semi-thin sections (0.5 -1 µm thick) were cut and stained with toluidine blue, until the fiducial marks (box) in the GCL appeared. The block was then trimmed and mounted in the SBFSEM microscope (GATAN/Zeiss, 3View). Serial sections were cut at 70 – 80 nm thickness and imaged at an x-y resolution of 6.0-8.0 nm. 3 x 3 tiles, each about 40 µm x 50 µm were obtained with an overlap of about 10%. The image stacks were concatenated and aligned using TrackEM (NIH). The MAC was traced using the tracing tools in Track EM.

Two additional SBFSEM datasets of fixed mouse retina spanning the IPL were used to explore glial ensheathment in more detail. The first block, k0725, covers a volume size of approximately 50 x 210 x 260 μm, with a voxel size of 13.2 x 13.2 x 26 nm (Ding et al., 2016). This resolution allows for the visualization of vesicles and chemical synapses, but does not allow for identification of gap junctions. The second block used, k0731, was of the approximate size 60 x 80 x 80 μm with a similar voxel size to the k0725 block. The preparation of k0731 block was optimized to preserve extracellular space within the retinal tissue, allowing for identification of gap junctions (Pallotto et al., 2015).

To begin reconstruction analysis, Müller cells were easily identified due to dense internal filament structures characteristic of Müller glia that are absent from other cell types in the IPL. From Müller cell reconstructions, neuronal ensheathments were identified and ensheathed cells were reconstructed, leading to the identification of two narrow-field amacrine cells whose morphology and synaptic organization matched that of the MAC. Inputs and outputs on these cells were then counted and categorized. Bipolar cell inputs were matched to a bank of previously reconstructed bipolar cells from the same block in order to identify bipolar cell type (Ding et al., 2016).

All cells from these two blocks were reconstructed and annotated using the electron microscopy analysis software Knossos (https://knossos.app/), and skeleton renderings for figures were created using the visualization software Paraview (Ahrens, 2005).

### Viral Tracing

All animal husbandry and procedures for these experiments were in accordance with the Longwood Medical Area Institutional Animal Care and Use Committee guidelines. Animals were kept and bred in biosafety containment level 1 conditions at all times, except during and after the VSV injections, which took place in biosafety level 2 conditions.

Neurod6-Cre (NEX-Cre) mice (a generous gift from Klaus Nave from University of Gottingen) were described previously (Goebbels et al., 2006). NEX-Cre mice were maintained on a C57BL/6 background, and crossed to conditional TVA-expressing (cTVA) mice generated either by the Saur Lab (Seidler et al., 2008) or by the Cepko Lab (Beier et al., 2013). During initial studies, a conditional tdTomato reporter mouse (The Jackson Laboratory, stock number: 007909, commonly known as Ai9) was also bred to the cTVA mice to obtain a Cre-driven tdTomato reporter and TVA receptor expression. Zero viral transmissions were observed from an infected RGC to an amacrine cell that was not tdTomato+, confirming our previous findings of the specificity of envA-TVA transmission (Beier et al., 2013).

As an alternate way to deliver TVA, we employed plasmids or AAV viruses that encode Cre-dependent TVA. Plasmids or AAVs were subretinally injected into conditional NEX-Cre mice at P0-1. Approximately 0.3μl of virus, AAV-flex-Tcb (2/8) (10^13^ gc/ml), or 0.3μl of either one of the plasmids, pAAV-flex-TC66t or pAAV-flex-Tcb (1μg/ul) (Miyamichi et al., 2013) were injected using a pulled angled glass pipette controlled by a Femtojet (Eppendorf) into the right eye. In the case of plasmids, an electric pulse was delivered right after the injections, in order to electroporate the construct (AAV preparation and electroporation strategies described previously in detail: (Matsuda and Cepko, 2007, Xiong et al., 2015). For the final analysis, data derived from the different methodologies employed for TVA delivery were combined.

In an effort to increase viral transmission efficiency, 11 out of 74 mice injected in the studies were bred to IFNαR^0/0^ mice (032045-JAX; MMRRC, (Drokhlyansky et al., 2017).

VSV construction and production strategies employed in this study were previously described (Beier et al., 2013). Briefly, a recombinant VSV with a chimeric glycoprotein (A/G) that consists of the extracellular and transmembrane domains of EnvA and cytoplasmic domain of rabies virus glycoprotein (RABV-G) was pseudotyped with RABV-G. All experiments were performed using mice of either sex after weaning, typically around 2-3 months of age.

The pseudotyped VSV(A/G)EnvA was injected unilaterally into the left dLGN, employing a stereotaxic instrument (Narishige International USA) and pulled capillary microdispensers (Drummond Scientific). Injection volume was 250-500nl (at a 100 nl/min rate, using an UltraMicroPump III (WPI). VSV(A/G)EnvA had a concentration of 1-2×10^8^ ffu/ml. Injection coordinates used were: A/P -2.5 from bregma, L/M 2, D/V -2.75.

After 2-4 days following VSV injections, mice were euthanized with CO_2_, and eyes enucleated. Corneas were removed with small scissors, lenses were removed, and retinae were isolated in cold PBS. Freshly dissected retinae were fixed in ice cold 4% formaldehyde for 30 minutes before 3x wash with PBS. Whole retinal cups were blocked in 5-6% donkey serum in PBS with 0.3% Triton X-100 (blocking solution) for 1 hour. Retinae were switched to the primary antibody in blocking solution (see below for concentrations) and incubated overnight at 4^0^C. The following day, retinae were washed 3x with PBS, switched to secondary antibody in blocking solution for 3 hours at room temperature, and washed again 3x with PBS (second wash included a DAPI co-stain). Finally, all retinae were mounted in Fluoromount-G (Southern Biotech).

<u>Primary antibodies used in these experiments were as follows:</u>

Rabbit anti-PPP1R17 (Sigma, Prestige Antibodies; HPA047819; 1:200)
Chick anti-GFP (Abcam; ab13970, 1:200)

<u>Secondary antibodies used in these experiments were as follows:</u>

Donkey anti-goat Alexa Fluor 647 (1:250, Jackson ImmunoResearch)
Donkey anti-rabbit Alexa Fluor 647 (1:250, Jackson ImmunoResearch)
Donkey anti-chick Alexa Fluor 488 (1:250, Jackson ImmunoResearch)

Infected cells in retinal whole mounts were then imaged using an inverted Zeiss LSM780 microscope (Carl Zeiss AG) with 405-nm, 488-nm, and 633-nm lasers and a 40x oil objective (Plan Apo 40x/1.3 Oil).

We focused on regions with sparse RGCs labelling in order to isolate and image individual RGCs. A total of 74 NEX-Cre mice carrying conditional TVA were injected into the LGN to generate the RGC dataset. Imaged RGCs were divided into two groups as either “connected” (with a virally labeled AC within their dendritic arbors) or “unconnected” (with no other virally infected ACs within their dendritic arbors). RGC images were analyzed using Fiji for stratification depth and dendritic arbor area. The following criteria were used to categorize the RGCs: dendritic diameter, depth of arborization in the IPL, and general appearance of dendritic morphology, such as symmetry and density. Several manuscripts on mouse RGC types were cross-referenced for the final morphotype decision (Jacoby and Schwartz, 2017, Sun et al., 2002, Volgyi et al., 2009). The RGCs with overlapping dendritic arbors were not included in the analysis as they could not be judged reliably in terms of their morphologic features.

### Fluorescent In Situ Hybridization (FISH)

Probe generation and FISH protocol were previously described (Shekhar et al., 2016). Briefly, cDNA was derived from 5 week-old C57BL/6J retina (Jackson, 000664) following RNA extraction (RNAqueous Kit, Ambion, AM1912) and reverse transcription with Superscript III (ThermoFisher, 18080051). In order to generate the GlyT1 antisense probe, PCR was performed using the following primer sequences: Forward: aagggatgttgaatggtgctgt, Reverse: <u>gaaattaatacgactcactataggg</u>ccagcaagatgagcatgaagaa. Reverse primer starts with a T7 sequence adaptor to allow *in vitro* transcription (underlined). We synthesized the probe using the DIG RNA Labeling Mix (Roche, 11277073910).

Tissue samples were prepared according to the methods described in (Shekhar et al., 2016), with minor modifications. FISH was performed on Dact2-GFP retina sections. Retinae were dissected quickly and fixed in 4% PFA in PBS at room temperature for 25 minutes and washed with PBS (2x, 5 minutes each). Then, the retinae were passed through a sucrose gradient until they sank in cold 30% sucrose in PBS. Retinae were transferred to 1:1 30% sucrose:OCT and incubated for 1 hour in a cold room with gentle shaking. Retinae were embedded in the 1:1 30% sucrose:OCT solution for sectioning. 50μm-thick sections were adhered to Fisher Superfrost Plus microscope slides coated with poly-d-lysine in 1x borate buffer, washed 2x with ddH_2_0 and dried completely. After the sections were air dried, they were treated with Proteinase K (1.5 μg/ml, NEB, P8107S), post-fixed in 4% PFA followed by acetic anhydride treatment. After probe hybridization, the GlyT1 probe was detected using anti-DIG-HRP (1:750) and tyramide amplification (Perkin Elmer TSA Cy3). This was followed by the immunohistochemistry protocol to amplify the GFP signal in Dact2-GFP retinas. Chick anti-GFP (Abcam; ab13970, 1:200) and donkey anti-chick Alexa Fluor 488 (1:250, Jackson ImmunoResearch) were used, as described in the viral tracing section of the Materials and Methods.

## Results

A recent inventory of the neuronal makeup of the mouse IPL using serial block-face EM reconstructions (Helmstaedter et al., 2013) revealed >45 morphologically distinct types of amacrine cells. When ranked by density (# of cells per square millimeter of retina), the majority of the well-studied amacrine cells, including those involved in night vision (AII, A17) and the encoding of directed motion (On and Off starburst amacrine cells, or SACs) are among the top ten (Figure 2A). Also, near the top of this list are several amacrine types for which very little is known. More than half of the high-density amacrine types display narrow-field morphologies, which according to previous work, suggests that they use glycine as a neurotransmitter (Figure 2A,B; (Menger et al., 1998, Pourcho and Goebel, 1983), but this has not been confirmed. We focused on the second most densely distributed amacrine type, initially termed AC51-70 (Helmstaedter et al., 2013), and now referred to as the MAC. These cells are easily recognized by their unique morphology, as they are the only narrow-field amacrine cells that stratify exclusively in the ‘ON’ layers of the IPL (corresponding to the inner half of the IPL; Figure 1), and have only 1-2 neuritic appendages that connect the synaptic arbor to the soma. Their functional characteristics and circuit wiring, however, remain largely unknown (Pang et al., 2012). Cells with similar morphology also have been observed in the primate retina (Polyak, 1941), but as with mouse, we know little about them. Here we combine several techniques to explore the synaptic connectivity and physiology of MACs in the mouse retina.

**Figure 2.**
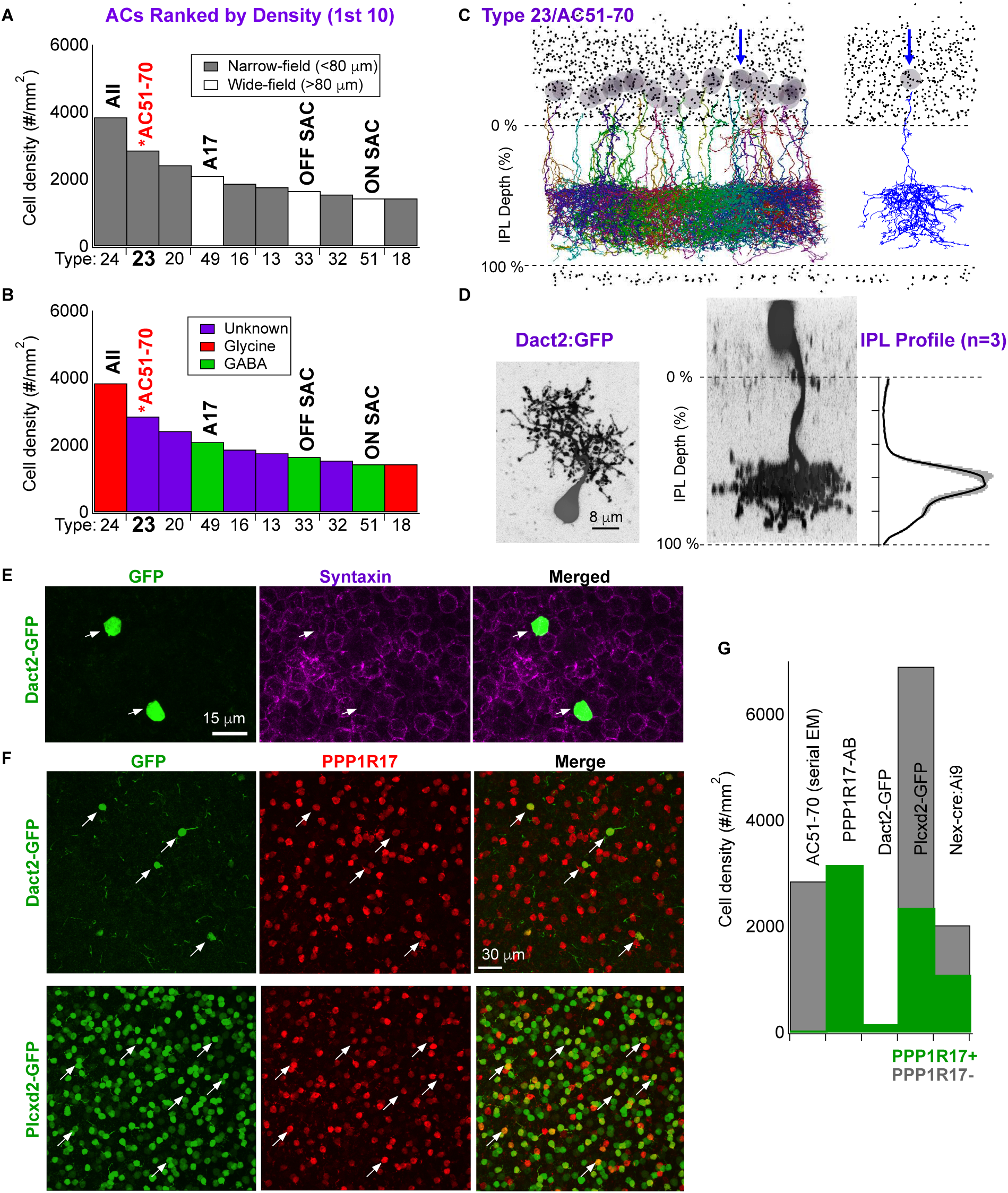
Identification of a high-density, narrow field amacrine cell, the MAC (aka AC51-70 and type 23). A) The ten amacrine cell types with the highest spatial density in the mouse retina ranked according to density (Helmstaedter et al., 2013). B) The top ten amacrine cell types (as in color coded according to what is known regarding their inhibitory neurotransmitters (i.e. GABAergic, Glycinergic, unknown). C) *left* Reconstructions of all MACs (AC51-70s) adapted from Helmstaedter et al., 2013. *right* Example of a single MAC; the blue arrow points to the corresponding cell in the population. D) MACs are uniquely and sparsely labelled in the *Dact2-GFP* line. Top and side views of a MAC labelled with GFP in the *Dact2-GFP* mouse line. Three isolated and strongly-labelled MACs (from the *Dact2-GFP* line) were used to assess stratification within the IPL (thick line represents the mean, shaded region represents SEM). E) Anti-syntaxin, an antibody specific to amacrine cells, labeled GFP+ cells in the *Dact2-GFP* line. F-G) Anti-PPP1R17 labeled amacrines at a density similar to that of MACs found in serial EM literature, furthermore all GFP+ cells in the *Dact2-GFP* line were PPP1R17+. G) Other mouse lines used in this study, such as *Plcxd2-GFP*, *NEX-Cre-Tdt*, and *Dact2-cre-Tdt-ChR2*, labeled various fractions of the PPP1R17+ cells (MACs).

We identified a mouse line (*Dact2-GFP*) in the GENSAT (www.gensat.org) database that selectively labels a sparse but homogeneous population of amacrine cells with morphology matching the AC51-70 from Helmstaedter et al., 2013 (Figure 2C,D). Closer inspection of the cells’ morphology revealed a monopolar neuron with a small bushy arbor restricted primarily to the inner (i.e. ON) layers of the IPL, and immunohistochemical (IHC) experiments conducted on this line showed that this cell type expresses the amacrine marker, syntaxin (Figure 2E; images acquired at the amacrine cell body level of the INL; (Voinescu et al., 2009)). Additionally, we mined the single-cell transcriptomic database for the mouse retina from Macosko et al. with the purpose of finding markers for the MAC (Macosko et al., 2015). In their paper they identify a cluster (cluster 20) which was fairly unique among retinal cells in its expression of the protein phosphatase regulatory subunit, PPP1R17. They determined that an antibody to PPP1R17 labels a population of narrow-field amacrine cells whose neurotransmitter and physiology were unknown. We found that an antibody directed to the phosphatase, PPP1R17, labeled 100% of the GFP+ cells in the *Dact2-GFP* line (Figure 2F,G), suggesting that it provides for an effective label of MACs. Furthermore, anti-PPP1R17 labelled cell bodies in the INL at a similar density to that reported for the AC51-70 in the Helmstaedter et al. dataset (Helmstaedter et al., 2013), Figure 2G) suggesting that this antibody might selectively label the entire population of AC51-70s. We later use this antibody to identify MACs in the absence of a genetically-encoded fluorescent label.

### MACs make connections with neurons and Müller glia

We targeted a GFP+ cell (i.e. MAC) in the *Dact2-GFP* line for ultrastructural analysis by combining serial block-face electron microscopy with the near-infrared branding (NIRB) technique (Bishop et al., 2011, Della Santina et al., 2016). Reconstruction of this MAC revealed a total of 26 synaptic inputs and 20 synaptic outputs, as indicated by clusters of vesicles adjacent to electron-dense regions of membrane (Figure 3A; we return to the cell’s synaptic partners in a later section).

**Figure 3.**
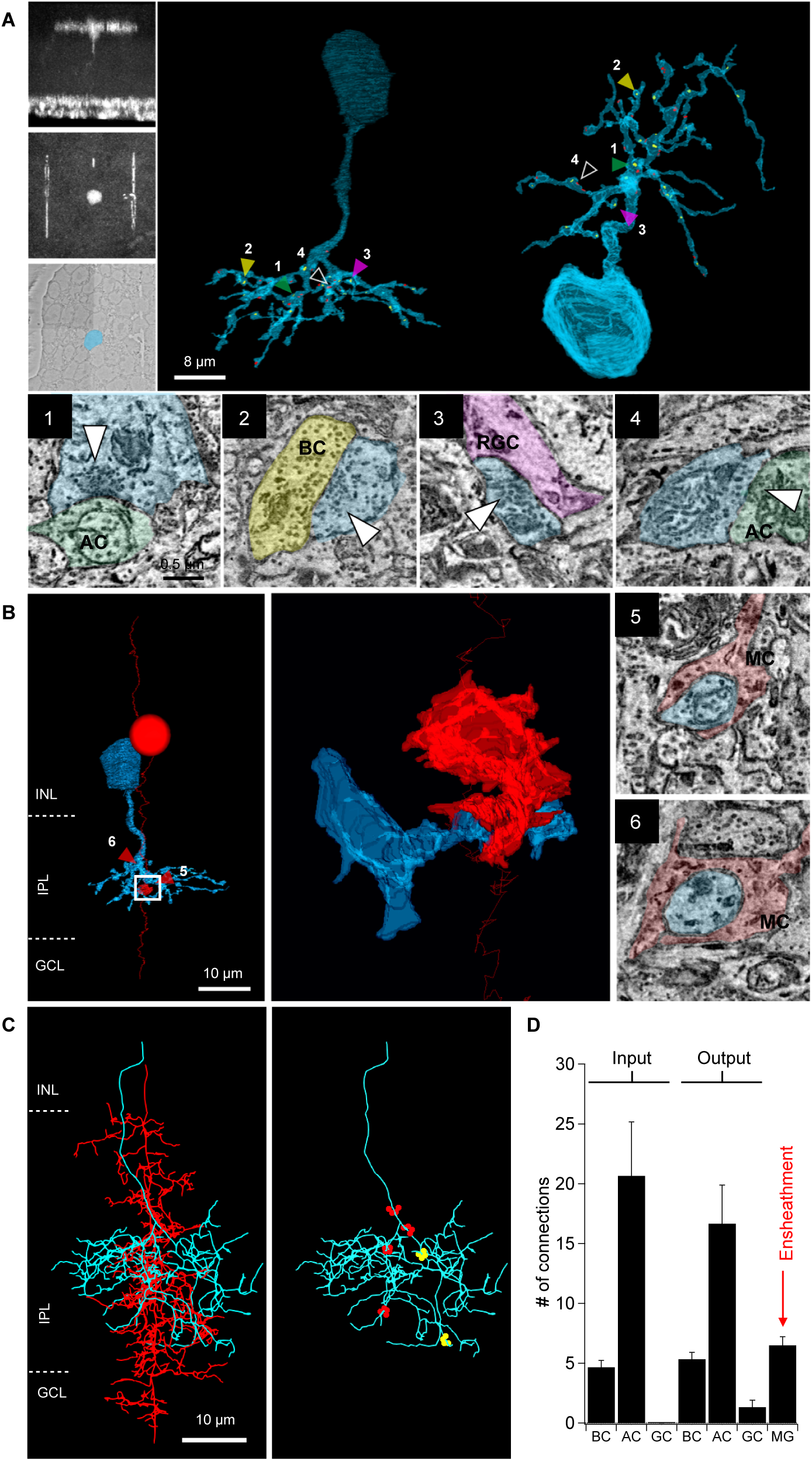
Serial block face EM reconstructions reveal canonical synaptic connectivity and Müller glia ensheathment. A) *left* NIRB marks were used to target a GFP+ cell in the *Dact2-GFP* line. *right* The resulting MAC reconstruction revealed conventional output synapses onto retinal ganglion cells (RGCs, 3), amacrine cells (ACs, 1), and bipolar cells (BCs, 2), and chemical synaptic inputs from bipolar (shown in Figure 5D) and amacrine cells (4). B) Reconstructions also revealed Müller glia ensheathment (red) of MAC processes (blue, 5, 6) that lacked obvious chemical synaptic structures. *left* Skeleton and partial reconstruction of a Müller cell that ensheathe the MAC at multiple locations. *Middle* Expanded view of the 3D Müller ensheathment from the white box in the left panel. *Right* Electron micrographs of the Müller ensheathments denoted in the left panel with numbers 5 and 6. C) Müller cells reconstructed in another, non-labelled EM block (Graydon et al., 2018) also ensheathed MACs (see Materials and Methods). *left* An example of a Müller cell reconstruction (red) with multiple ensheathments of an amacrine cell (blue) with morphology matching the MAC. *Middle* This particular MAC is ensheathed by the red Müller cell at 4 locations within its arbor (red ensheathments), and also with another Müller cell (not shown) at 2 additional locations (yellow ensheathments). Note that the cell bodies for these Müller glia and amacrine cells are missing from this thinner block which only contains a small fraction of the inner nuclear layer, where their cell bodies typically reside. D) Bar graph summarizing the ultrastructural analysis of 3 reconstructed MACs (Mean ± SD).

In addition to conventional synapses, the reconstruction revealed glial ensheathment on neuronal compartments that lacked obvious synaptic structures (Figure 3B). These glial ensheathments were found on the descending neuritic stalk and branches of the small bushy arbor of the MAC. We wondered if glia ensheathment was simply a consequence of high-density cell types with overlapping processes. However, glial ensheathment was rarely observed in complete reconstructions of two AII amacrine cells (i.e. the amacrine cell type with the highest retinal density; data not shown). To better determine the extent to which glial ensheathment is a true feature of MACs, we reconstructed Müller glia in a different block-face data set (Graydon et al., 2018) that did not have genetic labels or NIRB. Each Müller cell reconstruction (n=3) in this dataset exhibited multiple neuronal ensheathments throughout the IPL, and by reconstructing some of the ensheathed neurons (∼30) we found two amacrine cells which mirrored the morphology and synaptic organization of the MAC identified using the NIRB technique (Figure 3C). These data indicate that Müller ensheathment is a feature of MACs, but also suggest that some neurons, other than MACs, also are ensheathed by Müller glia. We later revisit the nature and specificity of neuronal ensheathments regarding retinal cell type.

We hypothesized that these ensheathments might enable interactions between neuronal and glia networks, and that gap junctions, which are not easily resolved in conventional serial block-face reconstructions (Pallotto et al., 2015), may play a role. To test for gap junctions between MACs and Müller glia, we injected gap junction permeable (neurobiotin) and impermeable (Lucifer yellow) tracer molecules into MACs (under light-adapted conditions) and analyzed the diffusion of these tracers into coupled cells using IHC (Mills and Massey, 1995, Pang et al., 2010, Pang et al., 2013, Vaney et al., 1998). These experiments were conducted on another transgenic mouse line (*Plcxd2-GFP,* Figure 2F) which had a much higher percentage of GFP+ MACs than the *Dact2-GFP* line (see Figure 2G for more information on overlap between PPP1R17 labeling and fluorescent protein expression in the various mouse lines used in this study). Tracer coupling experiments consistently (12 for 12) produced neurobiotin labeling in the Müller glia (recognized by their characteristic morphology) surrounding the injected MAC, but not neighboring MACs (Figure 4A). In contrast, Lucifer yellow labeling (included in a subset of injections; n=4) was constrained to the injected MAC (Figure 4A). To further confirm the identity of putative Müller glia, we also used IHC for CRALBP, a known marker for Müller glia (Bunt-Milam and Saari, 1983). Indeed, suspected Müller glia were immuno-reactive for CRALBP (data not shown). This experimental approach occasionally showed additional tracer-coupling to cone bipolar cells and wide field amacrine cell processes, but these coupling patterns tended to be variable across samples and were thus not pursued further.

**Figure 4.**
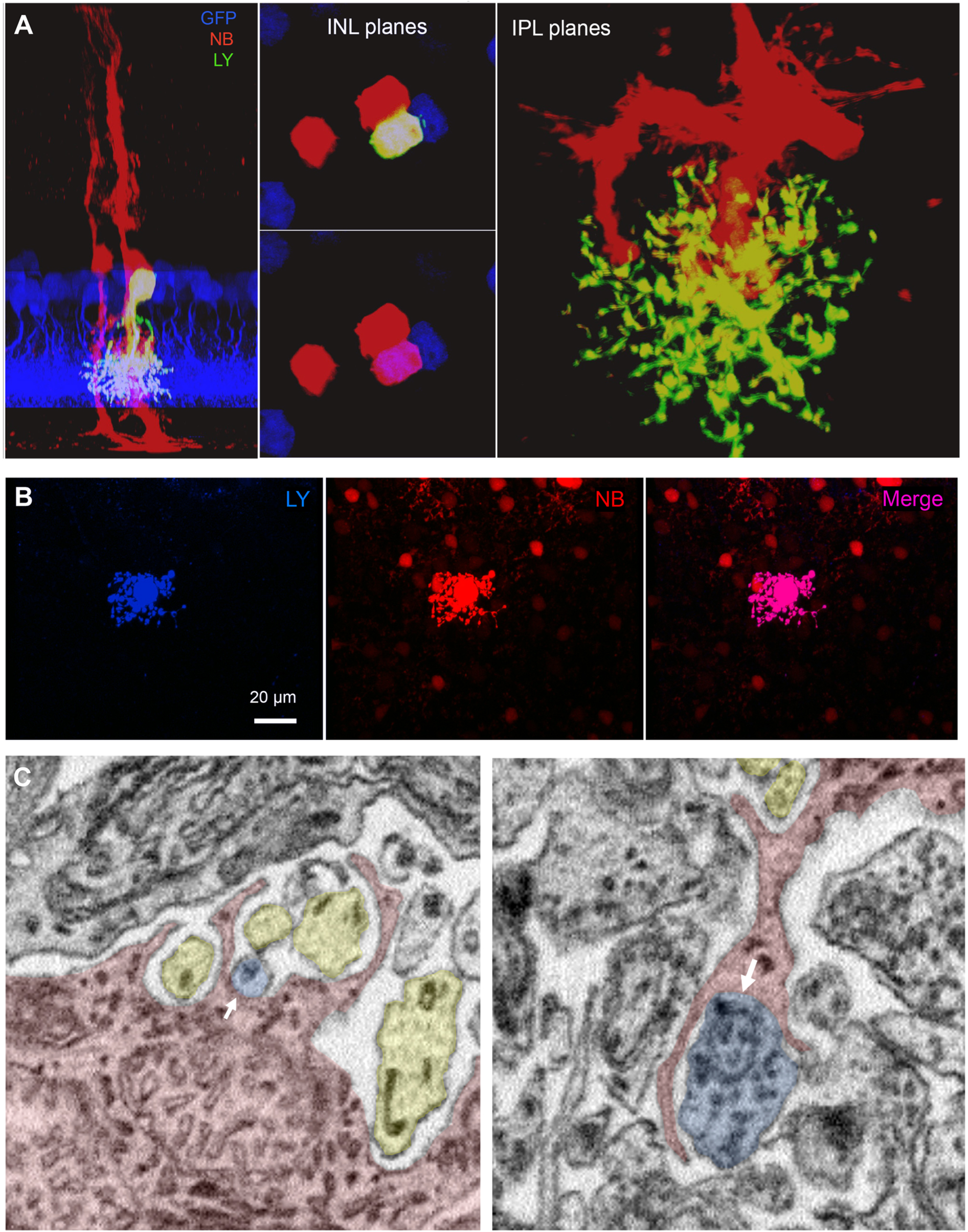
MACs are gap junction-coupled to Müller glia. A) A single MAC in the *Plcxd2-GFP* line was injected with gap junction-permeable (neurobiotin, NB) and -impermeable (Lucifer yellow, LY) tracer molecules and analyzed using IHC. Neurobiotin labelled nearby Müller glia, whereas Lucifer yellow was only observed in the injected MAC, indicating a selective transmission through gap junctions. B) A single AII amacrine cell injected and analyzed using the same tracer coupling method. Other neurons were labelled by neurobiotin, but Müller glia cells were not. C) Electron micrographs of contact (presumably gap junctions) between Müller glia (red) and amacrine cells (blue) in the IPL (block from Pallotto et al., 2015). Yellow shading denotes ensheathed neurons without obvious gap junction connectivity.

The observed coupling with Müller glia was not a feature common across amacrine cell types, as similar injections into AII amacrine cells (n=2) labelled On cone bipolar cells and a few AII amacrine cells but did not label Müller glia (Figure 4B; see also (Mills and Massey, 1995, Trexler et al., 2001)). We were unable to identify gap junctions in the EM datasets discussed thus far, so we turned to another published EM dataset where gap junctions are identifiable due to preservation of the extracellular space (Pallotto et al., 2015). Under normal conditions, nearly 20% of the brain’s volume consists of extracellular space, which creates a separation between the membranes of neighboring cells except when a synapse (chemical or electrical) is present.

Typical EM fixation protocols utilize hypoosmotic solutions that lead to a collapse of extracellular space, leaving gap junctions, which aren’t typically surrounded by clusters of intracellular vesicles, difficult to resolve. Although we were unable to find a complete MAC in this much smaller block (see Materials and Methods), partial reconstruction of a Müller glia cell (red) revealed that some neuronal ensheathments in the IPL contained small gap junctions with amacrine cells (blue), whereas other ensheathed cells—e.g. bipolar and ganglion cells—lacked gap junction connectivity (yellow; Figure 4C). These data indicate that only a fraction of a Müller cell’s neuronal ensheathments contain gap junctions, and that these connections seem to be made preferntially with amacrine cells.

### Physiological response to visual stimulation

The light encoding properties of MACs were investigated by targeting GFP+ cells for whole cell recordings using two-photon microscopy (980 nm) in dark-adapted *Plcxd2-GFP* retinas. We began by probing the receptive fields of MACs with brief presentations of spots of various sizes (50-1000 um) and contrasts (±100% on a 200 R*/rod/s background). Current clamp recordings revealed a clear center-surround receptive field (Figure 5A). MACs showed a strong preference for positive contrasts (i.e. light increments) regardless of spot size (Figure 5A-C), consistent with their stratification within the On region of the IPL (Figure 2D).

**Figure 5.**
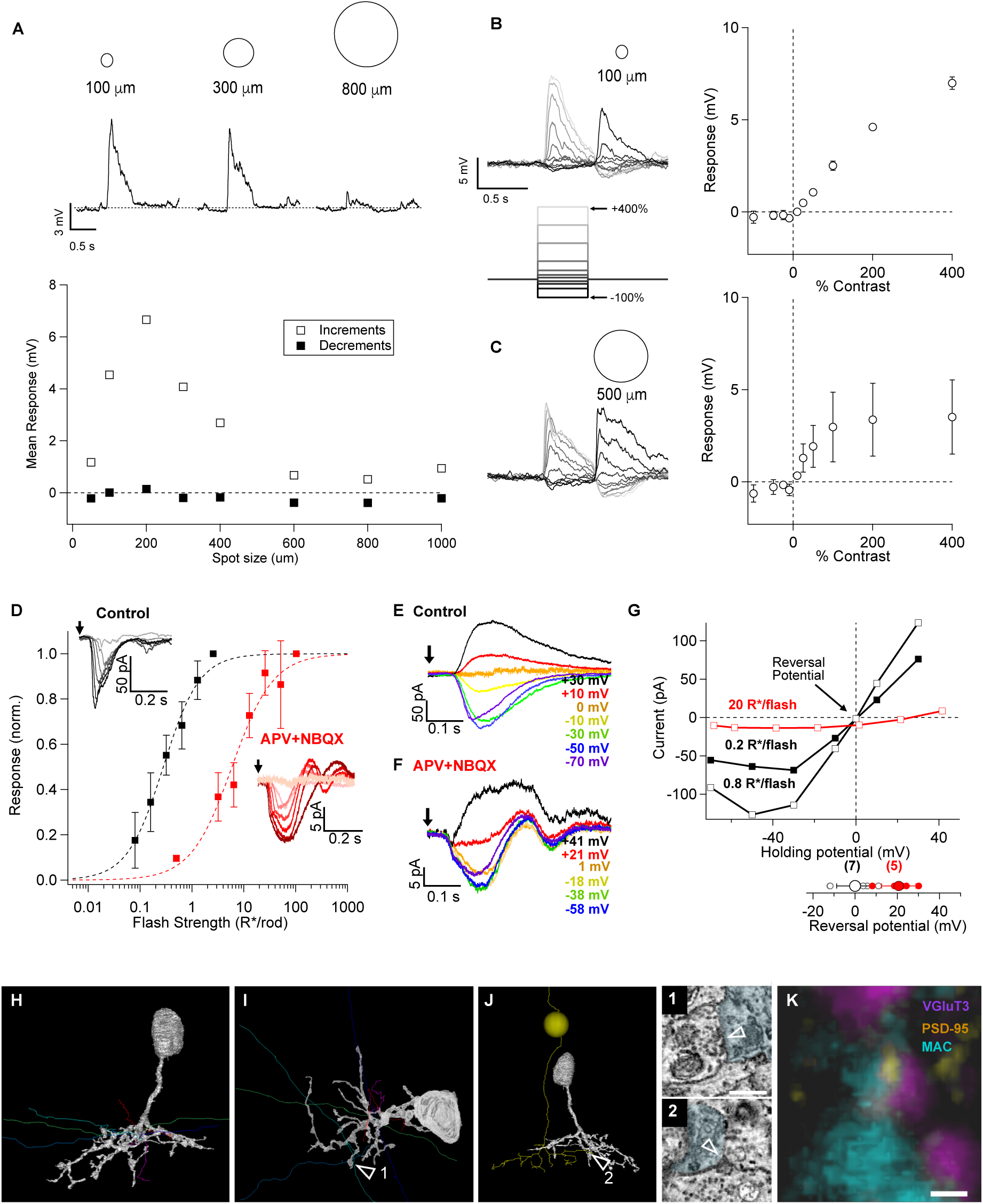
Visually-evoked signals and origins of excitatory synaptic input to the MAC. A) *top* Current clamp recording of visual responses to flashed spots (light increments) of increasing size. *Bottom* Response amplitudes plotted as a function of spot size for the example cell. Responses to light increments are denoted with open markers and responses to light decrements are denoted with closed markers. B) Responses to a 100 micron spot of various contrasts. (n=4) C) Responses to a 500 micron spot of various contrasts. (n=4) D) Voltage-clamp recordings (Vhold ∼ reversal potential for chloride) assessing flash sensitivity from darkness. Normalized response amplitudes are plotted as a function of flash strength. After control recordings APV and NBQX were bath applied to block NMDA and AMPA receptors, respectively. Responses to flashes *≤*1 R*/rod/flash were blocked by the cocktail, but further increasing the flash strength under these conditions revealed cocktail-insensitive responses. E,G) Dim flash responses in darkness reversed at the theoretical reversal potential for excitatory glutamate receptors (∼0 mV). F,G) APV- and NBQX-insensitive light responses emerged as flash strength was increased, and reversed at positive potentials indicating a lack of voltage-clamp. H-J) Cells providing synaptic input to MACs as observed with serial EM. H,I) Side view (H) and top view (I) of the reconstructions of presumed wide-field amacrine processes directly presynaptic to the MAC. J) Side view of a cone bipolar cell that makes two ribbon-type synaptic contacts with the reconstructed MAC. I_1_) Electron microscopy image of a conventional input synapse denoted by the arrow in I. J_2_) Electron microscopy image of a ribbon input synapse denoted by the arrow in J. K) Rare example of colocalization between fluorescently-tagged VGlut3 antibody (purple), YFP-tagged PSD-95 (yellow) and fluorescence from a single MAC (fluorescently-tagged streptavidin labeling of neurobiotin loaded through micropipette, blue).

We next explored the synaptic mechanisms mediating flash (10 ms flash, 500 um spot) sensitivity from darkness using the voltage-clamp recording configuration. At a holding potential of ∼−68 mV, MACs were sensitive to dim flashes that elicited ∼ 0.1 photoisomerizations per rod per flash (R*/rod/flash) and were largely saturated by flashes greater than 1 R*/rod/flash (Figure 5D). Flash strength was then held constant while varying the holding potential. These flash-evoked signals reversed near 0 mV (−0.05±3.3 mV, n=7, Figure 5E, G), the reversal potential for excitatory synaptic currents (e.g. mediated by AMPA or NMDA receptors). The ability to null current flow at this holding potential suggests that the cells primarily receive excitatory, not inhibitory, synaptic input under these conditions (Figure 5E). Furthermore, the current-to-voltage relationship for the synaptic inputs was j-shaped (Figure 5G), a characteristic of synaptic inputs mediated at least in part by NMDA receptors (Dingledine et al., 1999, Manookin et al., 2010). Responses to flashes producing 1 R*/rod or less were almost entirely abolished by a combination of NMDA and AMPA receptor antagonists (Figure 5D). These drugs likely alter upstream visual signals in addition to their action on MACs. However, increasing the flash strength (∼30-fold above the half-maximal flash in control) under these pharmacological conditions led to the recruitment of an additional response component with an I-V relationship that differed from control conditions in two ways (Figure 5D, F, G). First, the I-V relationship exhibited a much shallower slope than in control conditions, and second, the responses reversed at significantly more positive holding potentials (20.7±3.6 mV, n=5, p=0.002; Figure 5G). These responses are not easily explained by excitatory or inhibitory synaptic inputs, which have respective reversal potentials of 0 and −68 mV for our recording solutions. Instead, the presence of a weakly voltage-dependent inward current is consistent with gap-junction-mediated input from a retinal cell type with a higher visual threshold (Grimes et al., 2014, Ke et al., 2014, Murphy and Rieke, 2011, Trexler et al., 2005). Potential candidates include ON cone bipolar cells, which receive direct synaptic input from cone photoreceptors through APV+NBQX-insensitive mGluR6 receptors (Dunn and Wong, 2012, Masu et al., 1995, Slaughter and Miller, 1981), and Müller glia (Figure 4).

Which cells mediate excitatory synaptic input to MACs? Our EM reconstruction revealed both conventional (Figures 2A_4_ and 5J_1_) and ribbon-type synaptic input to MACs (Figure 5J_2_). This observation, along with a lack of functional inhibitory responses to the stimuli used here (Figure 5), led us to hypothesize that MACs receive glutamatergic input from bipolar cells and possibly a glutamatergic amacrine cell. Recent studies have shown that, whereas amacrine cells are generally considered to be inhibitory, one known amacrine type expresses vesicular glutamate transporter 3 (VGLUT3; (Haverkamp and Wassle, 2004, Johnson et al., 2004, Grimes et al., 2011)) and releases glutamate (Lee et al., 2014, Lee et al., 2016). Morphological assessment of stratification overlap in the central layers of the IPL suggests that VGLUT3 ACs could provide input to the MAC (Figure 2D). We tested for these connections using both anatomical and physiological approaches. For our anatomical approach, we crossed the *Plcxd2-GFP* line with a mouse line that expresses YFP tagged to the glutamate receptor scaffold PSD95 and labelled these retinas with the VGLUT3 antibody. PSD95 was rarely observed at appositions between VGLUT3 ACs and MACs (Figure 5K), suggesting that VGLUT3 ACs provide little or no input to MACs. For our physiological approach, paired recordings (n=6) between MACs and VGLUT3 ACs (in a *Plcxd2-GFP*x*VGLUT3-cre-Tdt* mouse line) were conducted, but the recordings failed to show obvious signal transmission between these two amacrine cell types.

We also used our serial EM datasets to help identify the cell types that are presynaptic to MACs. Upon closer inspection we determined that the majority of the conventional synapses come from narrowly stratified wide-field amacrine cells whose processes tended to pass straight through our relatively small EM volume (Figure 5H,I). One would predict that these wide field amacrines provide GABAergic input to MACs, but we found no evidence for light-evoked inhibition in our MAC recordings. The nature of the synaptic connections with these wide-field amacrine cells remains a mystery for future studies. Determination of bipolar cell identity was facilitated by using EM data from previously published work (Ding et al., 2016, Graydon et al., 2018). Reconstructions of the neurons providing ribbon-type input to the 2 MACs found in the non-NIRB block were successfully identified as type 5 and type 6 cone bipolar cells (see Materials and Methods).

### MACs make conventional glycinergic synapses

Across all reconstructed MACs we found an average of 23 conventional chemical output synapses onto amacrine, bipolar, and RGCs (Figure 3D). To determine the molecular identity of the neurotransmitter released at these conventional synapses we turned to immunohistochemistry.

Previous anatomical studies found a correlation between amacrine arbor size and neurotransmitter type; narrow field amacrine cells typically express glycinergic markers (e.g. GlyT1) whereas amacrine cells with wide arbors typically express GABAergic markers (Menger et al., 1998, Pourcho and Goebel, 1983). IHC assessment of the *Dact2-GFP* line showed that narrow-field MACs are immunoreactive for glycine (Figure 6 *top row*), but negative for other aspects of the molecular machinery commonly associated with inhibitory cells (GlyT1, GAD67; Figure 6 *middle and bottom rows*). This result is consistent with transcriptome sequencing data for PPP1R17-expressing amacrines in Macosko et al., 2015.

**Figure 6.**
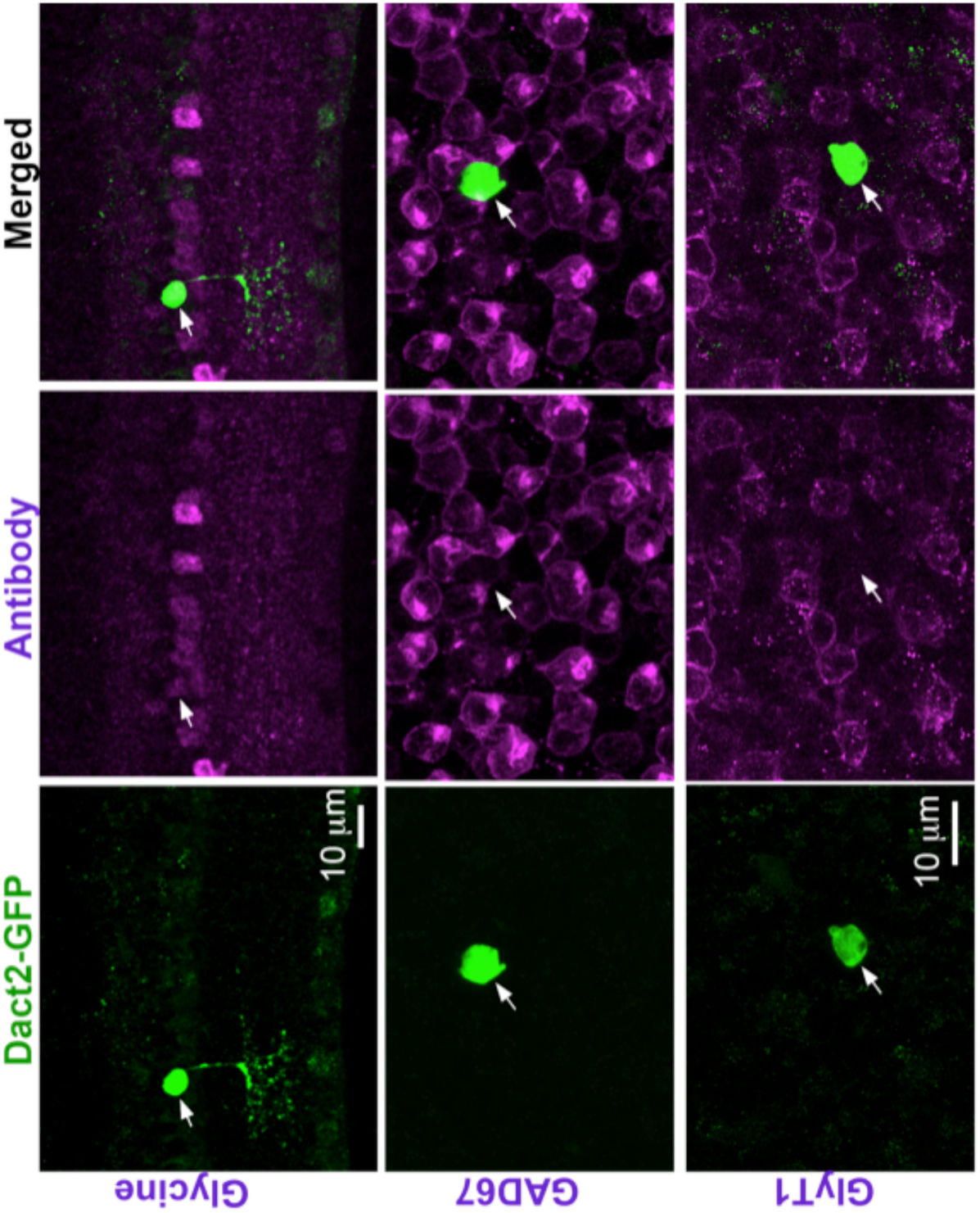
Immunolabeling experiments show that MACs express glycine, but antibodies against Gad67 and GlyT1 failed to label MACs. Antibody labelling for glycine (*Top row*), GAD67 (*middle row*), and GlyT1 (*bottom row*) in the retina of the *Dact2-GFP* line.

Together with the pattern of output synapses we observed in EM reconstructions, these data suggest that MACs release glycine onto multiple cell types in the IPL, including RGCs. To test for functional glycine release, we made recordings from RGCs in a mouse line that expressed ChR2 in a sparse subset of MACs (*Dact2-cre/Tdt-ChR2*). Unlike the *Dact2-GFP* line, which exclusively and sparsely labeled MACs, the *Dact2-cre/Tdt-ChR2* retinas also expressed tdTomato-tagged ChR2 in wide-field (presumably GABAergic) amacrines and RGCs. The wider repertoire of labelling with Tdt-ChR2 in the Dact2-cre line likely reflects the sensitivity and history of the activity of the Cre allele at this locus, but another cross (i.e. Dact2-cre x YFP-tagged ChR2) failed to label MACs entirely. We made whole-cell recordings from a random collection of RGCs in a flat-mount retinal preparation (Figure 7), and used a cocktail of APB, NBQX, APV and UBP to block glutamatergic synaptic transmission throughout the retina while leaving direct inhibitory synaptic transmission intact (Lee et al., 2014, Lee et al., 2016, Park et al., 2015, Tien et al., 2016). A spot of UV light was then delivered to the retina to test for ChR2-driven responses in RGCs. Only 1 of 48 RGCs tested showed a robust ChR2-evoked response. Synaptic currents recorded from this RGC (Figure 7A,B) reversed near the expected reversal potential for a Cl- mediated conductance and were completely blocked by the glycine receptor antagonist, strychnine (1 µM; Figure 7B). Subsequent imaging of the RGC and the ChR2-Tdt+ cells revealed a single MAC within the proximal arbor of a narrowly-stratified small arbor RGC that ramifies in the innermost layers of the IPL, with morphology similar to one of the W3 RGC types (i.e. Local Edge Detector; (Helmstaedter et al., 2013); Figure 7A). Other RGCs, like the On-Off DS cell, failed to respond to ChR2 activation (Figure 7C,D). The low success rate of these experiments likely reflects a combination of several features: sparse MAC labeling in the Dact2-cre/Tdt-ChR2 retina (most RGCs we tested had 0-1 MAC within their entire dendritic arbor), a low number of RGC contacts per MAC (Figure 2D) and a high degree of specificity in MAC connectivity with RGCs. The low success rate left us unable to establish RGC type-specific connections with any certainty. Nonetheless, this experimental approach provided an example of a MAC providing functional glycinergic output onto an RGC.

**Figure 7.**
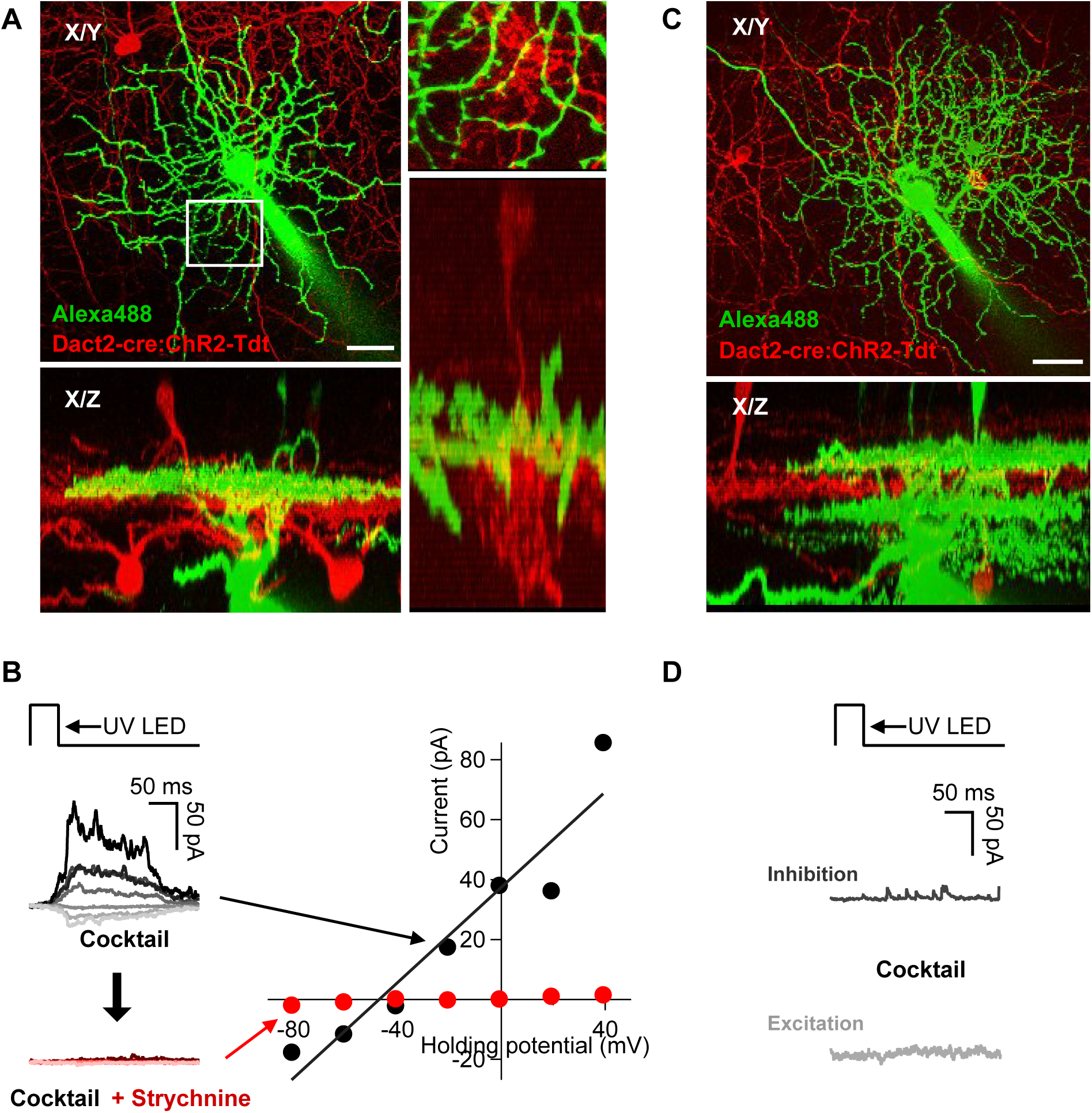
ChR2-evoked glycine release from the MAC. To test for glycinergic output from MACs, RGCs were randomly targeted in the *Dact2-Cre:ChR2-Tdt* line. Inhibitory input from ChR2-expressing cells was isolated with a pharmacological cocktail that blocks visual responses and glutamatergic synaptic transmission. A UV light step was then used to probe for positive connections, and Alexa 488 was included in the recording solution and imaged post-recording to aid in RGC identification. A) Example of a small arbor monostratified RGC (Alexa488, green) with one ChR2-Tdt(red)-labelled MAC within its dendritic arbor. B) ChR2-evoked responses recorded from the RGC in A for a range of holding potentials (i.e. −80 to +40mV). The amplitudes of the responses are plotted as a function of holding potential in the initial cocktail (gray traces), and in the presence of 1 uM strychnine (red traces). C-D) Example recording of an On-Off direction selective RGC in response to ChR2 activation in the *Dact2-Cre:ChR2-Tdt* line. C) Projections of an Alexa488-filled bistratified OODS RGC in the *Dact2-Cre:ChR2-Tdt* line. D) Voltage-clamp recordings of excitatory and inhibitory synaptic currents from the OODS RGC in C during ChR2 activation.

We also tested for glycine receptor expression at contacts between randomly-labeled RGCs and MACs using IHC. RGCs in the *Plcxd2-GFP* line were randomly injected with neurobiotin or labelled biolistically (see Materials and Methods). Retinas were then fixed and stained with antibodies against the alpha 1 subunit of the glycine receptor (GlyRα1; the GlyR isoform expressed on the dendrites of RGCs (Wassle et al., 1998)) and neurobiotin when relevant (Figure 8). This approach revealed GlyRα1 puncta at contacts between GFP+ cells and RGC types with various morphologies, including the suppressed-by-contrast RGC (Jacoby and Schwartz, 2018, Jacoby et al., 2015, Tien et al., 2016, Tien et al., 2015, Lee et al., 2016). By comparison, the OODS RGCs we encountered exhibited very little expression of GlyRα1 at appositions with GFP+ neuronal processes in the IPL (Figure 8B). Together these data argue that although MACs do not highly express GlyT1 (Figure 6), they do release glycine at conventional chemical synapses onto ganglion cells (Figures 7-8) and likely other neurons in the IPL (i.e. bipolar and amacrine cells, Figure 3D).

**Figure 8.**
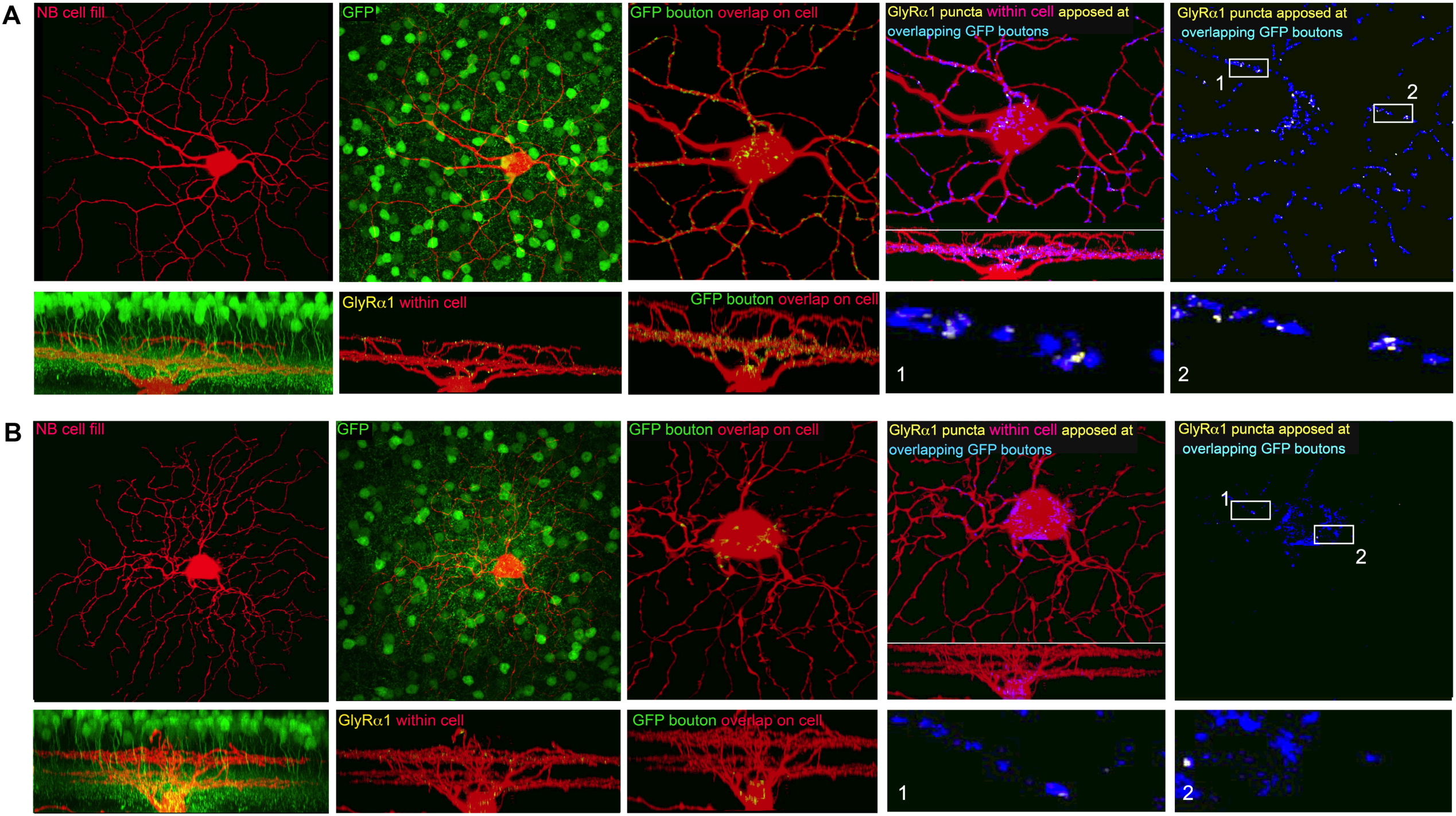
IHC identification of glycine receptors at putative contacts between MACs and RGCs in the Plcxd2-GFP line. RGCs were randomly injected with neurobiotin, prior to chemical fixation. GlyRα1 antibody and fluorescently-tagged streptavidin were used to label glycinergic synapses and neurobiotin-loaded RGCs, respectively. Overlap between neurobiotin and GFP signals was ascertained in 3D, together with the GlyRα1 signals within the dendritic processes. A) Example showing glycinergic contacts with a bistratified ganglion cell with morphology similar to that described for the suppressed-by-contrast RGC (Lee et al., 2016, Tien et al., 2015). B) Example of an On-Off direction selective ganglion cell with the same co-labelling protocol. GFP terminal appositions onto these cells rarely contained GlyRα1 puncta (in contrast to examples illustrated in A).

### MACs make synapses onto specific RGC types

We employed a viral tracing technique to more broadly assess MAC connectivity with RGCs (Figure 9). For this purpose, we used a mouse line with Cre expression in a more substantial number of MACs (NEX-Cre; (Kay et al., 2011)). This line also drives Cre expression in other amacrine cell types; thus, visual identification of the MAC’s unique morphology and/or the PPP1R17 antibody (Figure 2) was used to determine whether the connected amacrine cells were indeed MACs. We used a variant of VSV that expresses the avian EnvA glycoprotein, VSV-A/G(EnvA), to specify entry into cells that express TVA, the receptor for EnvA (Beier et al., 2013). The NEX-Cre mouse was used to specify the expression of TVA, either by crossing to a floxed-TVA mouse (Beier et al., 2013, Seidler et al., 2008), or by introduction of a flexed-TVA-mCherry construct (subretinal electroporation of the plasmid or subretinal delivery of an AAV vector carrying the flex-TVA (Miyamichi et al., 2013)). VSV was pseudotyped with the rabies virus glycoprotein (RABV-G) to allow the injected virus to be taken up by any cell at the inoculation site. Pseudotyped VSV-A/G(EnvA) was injected into a retinorecipient brain area (the lateral geniculate nucleus), where it was taken up by RGC axon terminals (Figure 9A). Retrogradely transported VSV-A/G(EnvA) replicates once it is in the cell body of a RGC, and then spreads transynaptically from the RGC to amacrine cells that express TVA. This method allowed us to probe many RGC types for connectivity to MACs. Animals were sacrificed at 2-4 days post VSV injection (dpi) to allow for transmission and expression of GFP from VSV-A/G(EnvA). Subsequent confocal imaging of these retinae showed that RGCs connected to MACs were enriched for several RGC types with distinct, known morphologies (Figure 9B,C,E): RGCs with small arbors (e.g. HD1, HD2 and W3;(Jacoby and Schwartz, 2017, Kim and Kerschensteiner, 2017); Figure 9B,E), 2) RGCs with highly asymmetric dendrites (e.g. JamB, F-mini;(Kim et al., 2008, Rousso et al., 2016); Figure 9C,E), and 3) RGCs that are asymmetrically bi-stratified (e.g. Suppressed-by-Contrast; (Jacoby et al., 2015, Tien et al., 2015); Figure 9E)). Although many different RGC types were labelled with this approach, all On-Off direction-selective (OODS) RGCs, a well-known RGC type that co-stratifies with the MAC, we encountered (n=10) lacked trans-synaptically-labelled MACs (Figure 9D). This lack of apparent connectivity between MACs and OODS RGCs is further supported by our IHC experiments that showed a lack of co-localization between NB-injected OODS RGCs, GlyRα1 puncta and GFP-labelled amacrine cells in the *Plcxd2-GFP* line (n=2; Figure 8B). Furthermore, and in contrast to our results, Beier et al., employing the same viral transmission strategy, found that when SACs expressed TVA, they were efficiently labelled by transmission of this same virus from OODS cells. Their data set showed a complete lack of connectivity between SACs and small arbor RGCs, despite the fact that small arbor cells were abundantly infected from their LGN viral injections (Beier et al., 2013). Taken together, the data in Figures 6-9 argue that MACs provide feedforward inhibition onto specific RGC types by way of conventional glycinergic synaptic transmission, while avoiding others (e.g. OODS).

**Figure 9.**
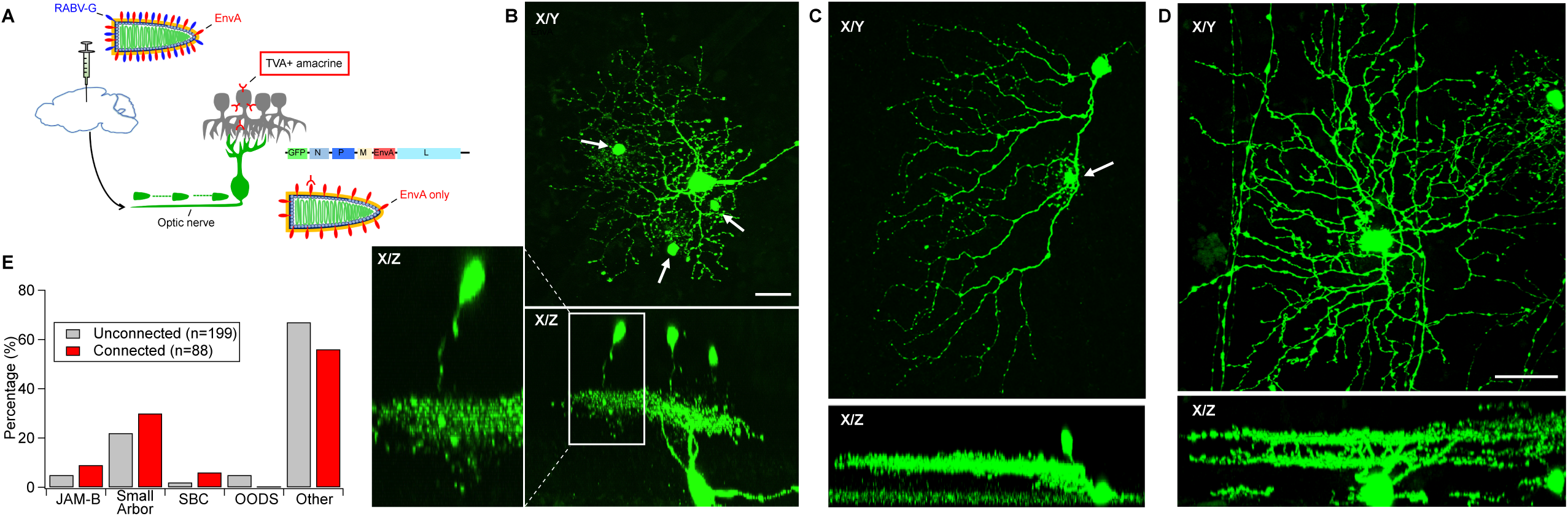
Assessment of MAC->RGC connectivity using viral tracing. A) A version of VSV that can infect virtually any cell type was injected into the LGN of mice. This form of VSV has the rabies G protein on its surface and can be taken up by axonal terminals and can travel retrogradely to a cell body. In the cell body, the virus can replicate and spread from an initially infected RGC. However, as the viral genome only encodes the EnvA glycoprotein, the virus can only transmit to cells expressing the TVA receptor. TVA expression was driven in these mice following cre-dependent recombination of a TVA allele in *NEX-Cre* mice, hence leading to TVA expression in a subset of MACs. Many RGCs were infected by these injections, but a large number of these RGCs did not transmit to any amacrine cell nearby (“unconnected” RGCs). RGCs with GFP-expressing amacrine cells within their dendritic arbors indicate a potential synaptic connection between the two cell types and were identified as “connected with MACs” RGCs. (B-D.) Views of RGCs and amacrine cells taken from the viewpoint of the X-Y plane and the Z plane. B) Example of a putative W3 RGC with multiple virally-connected ACs (arrows). Note co-stratification seen in the X-Y plane C) Example of a Jamb RGC (arrow) with one virally connected AC. D) Example of a virally-infected, unconnected OODS RGC (i.e. no connected amacrine cells). E) Quantitative assessment of the connected and unconnected RGCs from all injections.

## Discussion

Here we present our initial findings regarding the physiology and synaptic connectivity of the MAC, a recently discovered amacrine cell type that ranks second in amacrine cell density (i.e. cells per mm^2^) in the mouse retina. Insights from our experiments dramatically expand our knowledge of this high-density amacrine cell, but several interesting features remain to be resolved in future studies. It is our hope that progress on these questions will be facilitated by the mouse lines and antibody labels presented here (Figure 2).

One of these interesting features is the lack of traditional GABAergic and glycinergic markers (e.g. GAD67, and GlyT1) when assessed with IHC, which label most amacrine types. This observation is consistent with published RNA sequencing data on PPP1R17+ ACs (Macosko et al., 2015) and suggests that MACs are members of the newly discovered amacrine subpopulation originally termed non-GABAergic, non-glycinergic (nGnG) amacrine cells (Kay et al., 2011). However, our data indicate that these cells do release glycine via conventional synaptic transmission. We also note that our *in situ* hybridization did show *low* levels of GlyT1 within these amacrine (Figure 10). Thus, some apparent nGnG amacrine cells may release glycine despite low levels of expression of the commonly associated synaptic and transporter genes/proteins. It remains unclear how MACs acquire the appropriate levels of glycine, but previous work has shown that glycine can pass through gap junctions (Vaney et al., 1998), and that Müller glia, which we show are gap junctionally-coupled to MACs, take up extracellular glycine using GlyT1 (Hosoya et al., 2010).

**Figure 10.**
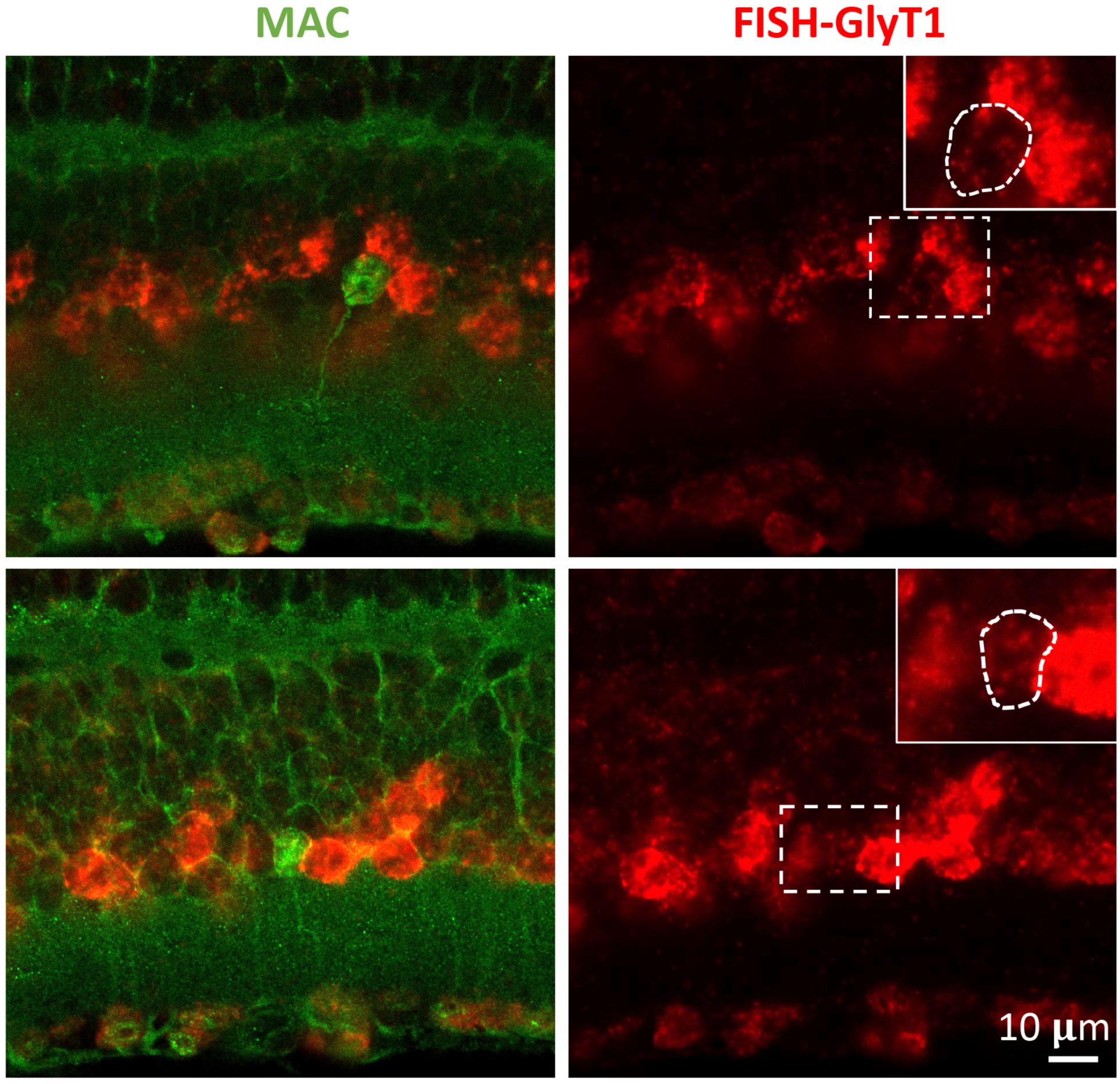
Low levels of GlyT1 transcript were detected in MAC. A FISH probe was employed against GlyT1 in Dact2-GFP retina sections. MAC cell body in green merged with the red GlyT1 signal on the left. GlyT1 signal visible in red on the right (dashed rectangle indicating where the MAC cell bodies are on the left, amplified on the upper right corner with the cell body outlined using dashed line). MAC is expressing GlyT1 at lower levels compared to the numerous glycinergic cells of the mouse retina, expressing GlyT1 strongly (bright red).

**Figure 11.**
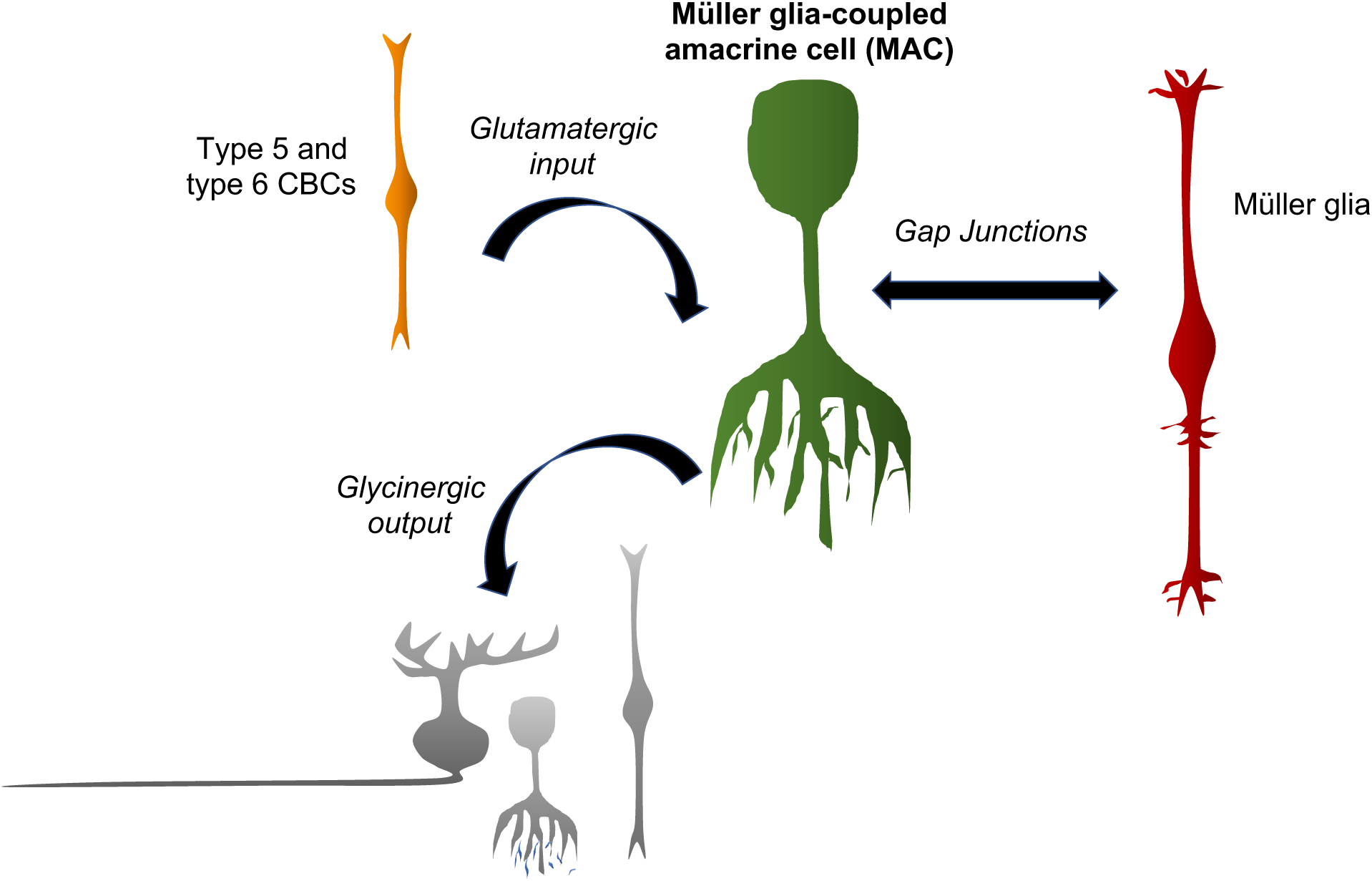
Summary of the MAC synaptic connectivity patterns found in this study. Our EM reconstructions confirmed that type 5 and 6 cone bipolar cells (CBCs) provide ribbon-type (glutamatergic) input to the MAC. EM reconstructions and tracer coupling experiments revealed gap junctions with Müller cells. Immunohistochemistry and ChR2 experiments indicate that MACs release glycine onto RGCs via conventional synapses, and EM reconstructions indicate that MACs make similar synapses onto bipolar and amacrine cells.

Our data also indicate that MACs receive light-evoked glutamatergic input but receive little or no inhibition in response to the visual stimuli utilized in our experiments (Figure 5). This physiology result is surprising given that, according to our EM reconstructions, the majority of input synapses to the MAC come from other amacrine cells (Figures 3D & 5H,I). One possibility is that these presynaptic amacrine cells release glutamate onto the MAC. Recent work has shown that one amacrine cell type expresses VGluT3 and releases glutamate (Lee et al., 2014, Lee et al., 2016), which led us to test for possible connections with the MAC. However, both paired physiology recordings and immunohistochemical analysis failed to show connectivity with the VGluT3 amacrine cell. Another possibility is that we have not yet found the visual stimulus that effectively drives these wide-field amacrines. And yet another possibility is that these interneurons provide tonic, but not phasic, inhibition to the MAC.

Lastly, we found that MACs are directly coupled to Müller glia. Direct coupling of neurons and glia is unusual (see (Alvarez-Maubecin et al., 2000, Pakhotin and Verkhratsky, 2005)) and has not previously been reported in the mammalian retina, but very few studies have tested this possibility directly. We speculate that MACs might play an important role in retinal functions involving neuron-glia interactions. For example, recent work (Biesecker et al., 2016) has revealed a neuron-glia interaction in the IPL that vasodilates retinal capillaries in response to increased neural activity (i.e. hyperemia). The identity of the neuron(s) mediating this function remains unknown.

While we do not yet fully understand the function of MACs, the results presented here concerning potential roles for MACs in visual processing and hyperemia are intriguing and should provide a basis for future studies.

## Notes

**Funding Sources** This work was supported by HHMI (C. Cepko, D. Göz Aytürk and F. Rieke), NIH grant (EY10699 to R. Wong), NIH Grant RO1 NS083848 (D. Göz Aytürk and C. Cepko), McPherson Eye Research Institute’s Rebecca Meyer Brown/Retina Research Foundation Professorship (M. Hoon), an unrestricted Grant from Research to Prevent Blindness, Inc. to UW Madison Department of Ophthalmology and Visual Sciences (M. Hoon), and by the NINDS Intramural Research Program (J. Diamond).

